# Multisystemic inflammatory disease in Pheasantshell (Unionidae, *Actinonaias pectorosa*) associated with *Yokenella regensburgei* infection at sites experiencing seasonal mass mortality events

**DOI:** 10.1101/2024.03.14.585088

**Authors:** Jeronimo G. Da Silva Neto, Rebecca H. Hardman, Augustin C. Engman, Gerald R. Dinkins, Timothy W. Lane, Michael M. Fry, Christian Rines, Amber Bisenieks, Sree Rajeev, Michelle M. Dennis

## Abstract

Freshwater mussels are integral components of riverine ecosystems, influencing water quality, nutrient cycling, and habitat characteristics. Enigmatic freshwater mussel declines, often characterized by sudden mass mortality events, pose significant challenges to conservation efforts. The Clinch River, a freshwater biodiversity hotspot, has experienced several enigmatic mass mortality events since 2016. Studies have reported bacteria associated with moribund Pheasantshell (*Actinonaias pectorosa*) during mortality events in the Clinch River, specifically *Yokenella regensburgei*. Despite reports of bacterial infection, little is known about their role as pathogens. Through a multiyear case-control study, combining in-situ experiments, field surveys, histology, bacterial isolation, and high-throughput sequencing, we assessed the role of bacteria in Pheasantshell (*Actinonais pectorosa*) mortality at two sites in the Clinch River. Between May 2021 and December 2023, we collected 29 wild moribund free-living *A. pectorosa* and 68 hatchery-reared *A. pectorosa* maintained in silos at the same sites and investigated differences in pathology and microbiology between groups. No silo mussels presented clinical signs of disease, or gross or microscopic lesions associated with pathological conditions leading to mortality. Our findings reveal a significant association between *Yokenella regensburgei* and severe multisystemic and multifocal infiltrative hemocytosis with necrosis, consistent with sepsis. Lesions associated with yokenellosis were of sufficient severity and physiological significance to explain mortality in infected hosts. Although our study does not explain the cause of these infections, it confirms that mussels at our study sites are ultimately dying from an infectious disease and that *Y. regensburgei* can be pathogenic in free-living mussels. Our results underscore the importance of considering bacterial diseases in wild mussel populations and emphasize the need for further research to elucidate the epidemiology and pathogenicity of *Y. regensburgei*. Overall, our study highlights the importance of integrated approaches combining pathology, microbiology, and epidemiology in freshwater mussel conservation efforts.

## Introduction

Freshwater mussels play an essential role in riverine ecosystem functioning. Mussels are benthic filter feeders that consume zooplankton, phytoplankton, and fine particulate matter from the water column, which influences water quality nutrient availability and cycling, algal and pollutant concentrations, and benthic productivity (Howard and Cuffey 2006; Vaughn 2018). In addition, mussels are considered habitat modifiers whose influence on the physical and chemical habitat characteristics of streams positively affect other freshwater fauna such as fish and macroinvertebrates (Sansom et al. 2018; DuBose 2020; Hopper et al. 2019).

Despite their ecological importance, mussel species richness in North America has been declining since the early 1900s at the alarming rate of 1.2% per decade (Ricciardi and Rasmussen 1999). Unionid mussels are arguably the most threatened group of freshwater fauna in North America with at least 27 of the 300 recognized species declared extinct and 65% of the remaining fauna considered imperiled (Ricciardi and Rasmussen 1999; IUCN 2017; Ferreira-Rodríguez 2019). Habitat alteration, loss, and degradation, declines in water quality, presence of invasive species, and disease are suspected of causing population declines (Haag and Williams 2014; Haag 2019). Although some population declines are attributed to degradation of habitat quality or exposure to pollutants, others have no clear cause (Haag 2019). These enigmatic declines have been reported across the globe for over 40 years, often in otherwise healthy streams with no apparent anthropogenic activity and where other aquatic fauna (e.g., fish, snails, and crayfish) appear unaffected (Haag 2019; Aldridge et al. 2023). Sudden mortality events, where hundreds or thousands of sick or dead mussels are observed, can be a component of enigmatic declines, and have been documented across south, central, and southern United States (Neves 1987; Haag 2019). These events can cause rapid declines in biodiversity and ecosystem services and can trigger long-term changes in community composition and widespread faunal collapse (DuBose et al. 2019b).

Despite the grave impacts, logistical and financial constraints often prevent the early detection of mass mortality events, which are typically reported incidentally to other monitoring efforts (Waller and Cope 2019). Therefore, they are seldom subjected to comprehensive diagnostic investigation and are poorly understood. The Clinch River watershed is a threatened freshwater mussel biodiversity hotspot. Located in the upper Tennessee River Basin of Virginia and Tennessee, the Clinch River contains 46 extant mussel species, over 20 of which are listed as federally endangered (Zipper et al. 2014; Jones et al. 2014). The Clinch River is a model system for exploring the potential causes of mass mortalities. Annual mussel mass mortality events have occurred in the river reach at the Tennessee/Virginia border since 2016, as highlighted by Richard (2018). In the Clinch River mortality events, large numbers of dead or moribund mussels have been observed between late September and November (Richard et al. 2021). Consistent with enigmatic declines observed elsewhere, this reach is considered healthy and only mussels seem to be impacted. First reported in Tennessee, mortality events have progressed upstream. At first, these mortality events appeared to affect all species (Richard 2018), but it became evident that Pheasantshell (*Actinonaias pectorosa*) exhibited greater mortality compared to other species (Richard et al. 2020). Pheasantshell are endemic to the Tennessee and Cumberland River drainages, and historically were especially abundant in the Clinch River compromising over 50% of the Clinch River’s mussel biomass (Jones et al. 2014). However, between 2016 – 2019, Pheasantshell populations declined 50 – 90% in some monitoring sites due to annual mass mortality events (Richard et al. 2020).

The disproportional loss of a single species and the upstream progression of the mortality events prompted research into the role of infectious microorganisms. Richard et al. (2020, 2021) reported the presence of a densovirus (Clinch densovirus1) and bacteria (i.e., *Salmonella enterica* and *Aeromonas salmonicida*) associated with moribund Pheasantshell collected from the Clinch River during mortality events. Similarly, Leis et al. (2019, 2023) reported high prevalence of bacteria (*Yokenella regensburgei*) associated with moribund mussels collected from the Clinch River in November during an active mortality event. However, the role of these viruses and bacteria in causing disease is uncertain, and there are no studies linking their detection to host pathology.

Here, we report the results of a multiyear case-control study conducted at two sites affected by mass mortality events in the Clinch River in Virginia and Tennessee. We used a combination of in-situ experiments, histology, conventional bacterial isolation, and high-throughput sequencing to assess the presence and etiological role of bacteria in the mortality of Pheasantshell. Our objective was to describe differences in pathology and associated bacteria between moribund and apparently healthy Pheasantshell. In addition, we provide a comprehensive morphological description of lesions associated with *Yokenella regensburgei* infection in order to establish a case definition for yokenellosis in Pheasantshell. Collectively, assessments of pathology and associated bacterial assemblages will help researchers and managers better understand how disease contributes to mass mortality events and overall mussel population declines in the Clinch River.

## Materials and methods

### Study Area

Our study area consisted of two sites in the Upper Clinch River watershed of Virginia and Tennessee. These sites were selected because 1) they have been affected by annual mortality events since 2016, 2) they have the last stronghold population of Pheasantshell, and 3) they are sources of broodstock for restoration efforts. Kyles Ford, Tennessee (36.564620, −83.041392), is located approximately 17 river miles downstream of Sycamore Island, Virginia (36.621061, −82.818235; Fig 1). Both sites are in the Upper Clinch River watershed (U.S. Geological Survey Hydrologic Unit Code 06010205), an area of approximately 1,960 *mi*^2^(5,100 *km*^2^) which encompasses approximately 500 linear miles (327 km) of the Clinch River. Land cover in the Upper Clinch HUC8 watershed is composed predominantly of deciduous, evergreen, or mixed forest (69%), followed by pasture/hay (17%), and urban development (8%; https://modelmywatershed.org/). Stream characteristics are similar at both sites, except Kyles Ford has a greater channel width (120m) and substrate more dominated by boulder and cobble dominant substrate compared to Sycamore Island which has a narrower channel width (60m) and a pebble/cobble dominant substrate. For field surveys, we selected smaller sampling units (20m x 60m) within the larger sites. We selected these survey units based on data previously gathered by numerous State and Federal agencies to encompass the greatest concentration of Pheasantshell in the area, thereby increasing our probability of encountering wild moribund mussels.

**Fig 1.**
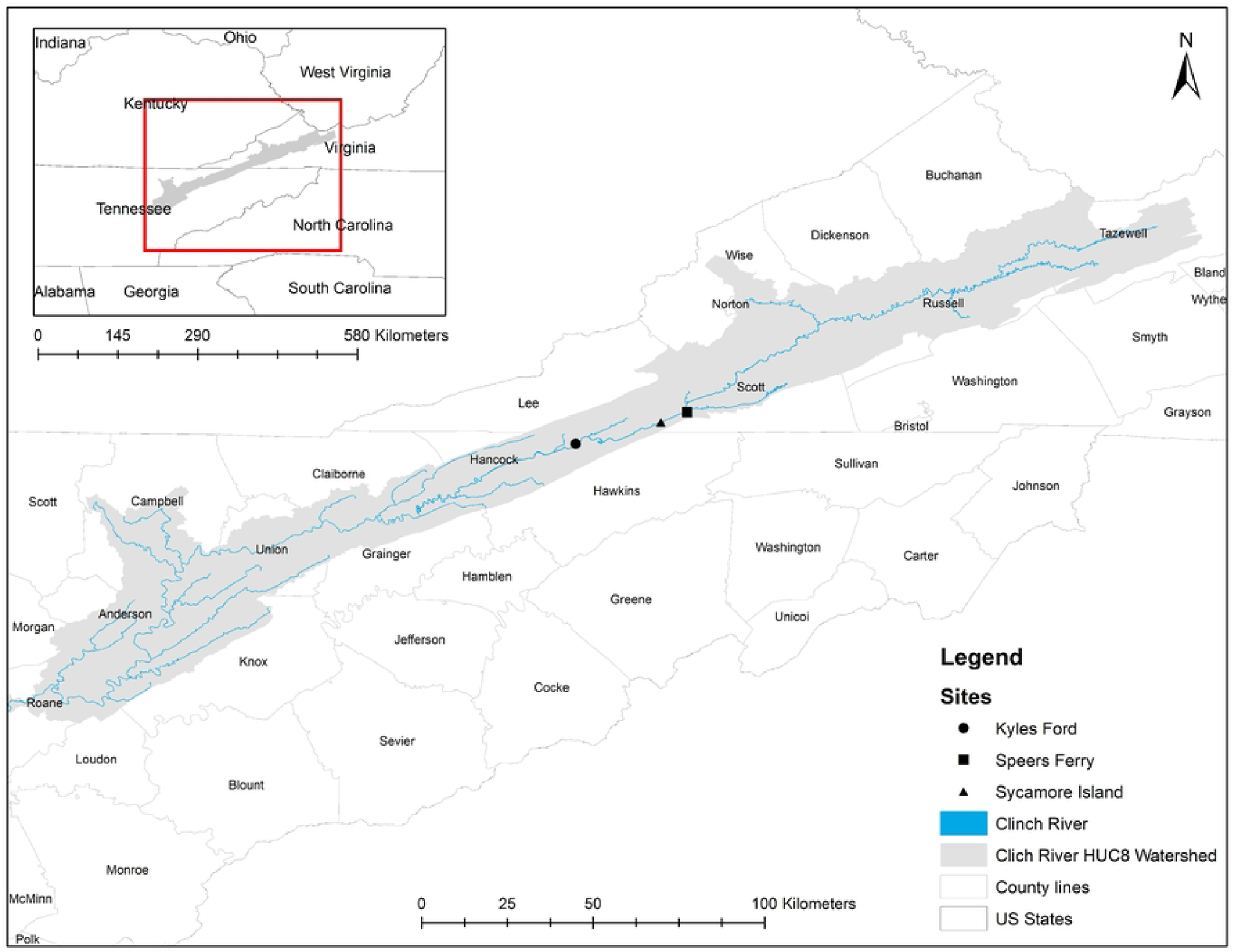
Map of study area. Map demonstrating the location of the Clinch River HUC8 watershed and the counties it overlaps. Our three sampling sites are in Scott County, Virginia, and Hancock County, Tennessee.

### Study design

We designed a control-case study that combined an in-situ experiment and field surveys. For the control group, we acquired 100 Pheasantshells (mean length = 13 mm) raised at the Virginia Department of Wildlife Resources’ Aquatic Wildlife Conservation Center. In May 2021 we deployed ten silos at Kyles Ford and ten silos at Sycamore Island (n = 20), and in each silo we placed five mussels. We randomly selected and removed five mussels from the silos at each site, once a month beginning in July 2021 to December 2021, and from September 2022 to December 2022. We did not collect mussels from silos between January 2022 and August 2022 because river discharge made sampling unsafe. If any mussels in silos showed signs of morbidity (i.e., gaping shell, flaccid foot, and diminished response to tactile stimulation), they would be harvested even if they were not randomly selected to be harvested. The case group was wild moribund Pheasantshells, collected during field surveys. We used a systematic sampling approach to sample wild moribund mussels occasionally between August 2021 – December 2021, and weekly between August 2022 – December 2022. We surveyed each sampling unit (20 m x 60 m) twice with two researchers for 30 minutes. During each survey, we positioned two researchers on opposite sides of the stream at the downstream end of the sampling unit. Starting from each side, researchers navigated to the midpoint of the river, walked upstream 1m and then proceeded away from the middle of the stream parallel to the initial direction. This pattern was repeated for 30 minutes until both researchers reached the upstream end of the sampling unit. Then, researchers would switch sides and conduct a second pass. During each pass, researchers collected any Pheasantshells that were displaying signs of morbidity (i.e., positioned unburied on the benthos, gaping shell, flaccid foot, and diminished response to tactile stimulation).

For each control or case mussel collected, we measured the long axis of each shell to the nearest millimeter using hand-held calipers and collected hemolymph in the field through aspiration of the anterior adductor muscle. Hemolymph samples were used for cytological assessment and bacterial 16S amplicon high-throughput sequencing analysis (16S sequencing hereafter). We collected 50 – 400 µL of hemolymph depending on mussel size. For each hemolymph sample, we aliquoted 50 µL into 1.5 mL sterile microcentrifuge tubes for 16S sequencing, 100 µL into 1.5 mL sterile microcentrifuge tubes pre-filled with 100 µL of L-cystine for cytological assessment and stored any remaining hemolymph into a 1.5mL sterile microcentrifuge tube. Hemolymph samples for 16S sequencing were transported in dry ice and then placed in a −80 °C freezer for future processing. Mussels were transported to the University of Tennessee Johnson Animal Research and Teaching Unit (JARTU) in coolers containing river water and aerators and placed in tanks with chlorine-free water to purge for 24h, which facilitated histological processing.

### Post-mortem examination and histopathology

After 24h we conducted autopsies on each mussel and prepared samples for histology. During autopsies, we opened each mussel by severing the adductor muscles. We used sterile equipment to remove a portion of the digestive gland for bacterial 16S sequencing and fixed the remaining tissue in 10% neutral-buffered formalin. Digestive gland samples collected for 16S sequencing were stored in a - 80 °C freezer until processing. We allowed tissue to be fixed for at least 48h, trimmed representative sections of all organs using a scalpel, and placed them in tissue cassettes which were submitted to the University of Tennessee College of Veterinary Medicine (UTCVM) Histology Laboratory for processing. Paraffin blocks were sliced in 5 µm sections and stained with hematoxylin and eosin (H&E). When indicated by pathology, slides were stained with a Hucker-Conn gram stain to assess bacteria presence associated with lesions. We used a chi-square test of independence to assess the association between microscopic lesions observed across all sites and clinical presentation (case vs control). The significance level for the analysis was set at alpha = 0.05.

### Hemolymph analysis

Once mussels were harvested, hemolymph was collected by aspiration of the anterior adductor mussel as previously described (Gustafson et al. 2005a) and was transported chilled in microcentrifuge tubes. We supplemented hemolymph with L-cysteine, stained hemocytes with trypan blue and used a hemocytometer to count hemocytes (Burkhard et al. 2009). For hemocytologic examination, we prepared cytocentrifuge smears stained with Wright’s Giemsa stain and examined each slide to assess hemocyte type and morphology. All hemolymph assessment was conducted at the UTCVM Clinical Pathology Laboratory.

### Bacterial 16S amplicon high-throughput sequencing

To determine bacterial taxa associated with observed disease, we conducted a comparative analysis of bacterial assemblage using 16S sequencing on hemolymph collected in the field and digestive gland collected during autopsies. All samples were immediately placed on dry ice or in the freezer after collection and stored at −80 °C until processing. We extracted DNA from samples using ZymoBIOMICS^TM^ DNA Miniprep Kits (Zymo Research Corporation, Irvine, CA, USA). We sent DNA aliquots to the University of Tennessee Genomics Core for 16S rDNA amplicon library preparation of a 250 bp section of the bacterial 16S v4 region and high-throughput sequencing on the MiSeq platform (v2 2 x 250; Illumina, Inc). For quality control for each sequencing run we had at least one extraction control, PCR blank, and extraction community standard (product # D6300, ZYMO Research Corporation).

Bioinformatic preparation of the resulting sequence data was performed in the program MOTHUR (Schloss et al. 2009) following the MiSeq standard operating procedure using operational taxonomic units (OTUs). OTUs were given taxonomic classification to genus based on the silva rdp reference database. We performed all subsequent cleanup and analyses using RStudio v1.2.5019 (R Core Team 2021). To evaluate only top contributing OTUs only from the abundance matrix, we removed OTUs with less than 500 total sequence reads across all samples. We also removed samples with less than 200 sequences based on rarefaction curves performed in package vegan (Oksanen et al. 2015). To compare relative OTU abundance, we used a 100-sequence subsample from all samples. Because of low sequence yield and low sample size for the majority of hemolymph samples they were removed from statistical analysis but were still included in plots for visual comparison. For digestive gland bacterial assemblages, we compared differences in presence of potential bacterial pathogens based on disease status (e.g., case vs control), and presence/absence of hemocytic nodulation and necrosis associated with gram-negative bacilli noted histologically. This lesion was absent in the control group. Therefore, we had three analysis groups: Control-Absent, Case-Absent, and Case-Present. We performed permutational multivariate analysis of variance (PERMANOVAs) to compare bacterial assemblages between groups via the ‘adonis2’ function in vegan and ‘pairwise.adonis’ function in package pairwiseAdonis (Arbizu 2017) using both Sorenson Index and Morisita-Horn dissimilarity matrices previously created from an average of 1000 iterations of 100-sequence subsamples. We graphically displayed community differences between groups of both hemolymph and digestive gland samples using non-numeric multidimensional scaling (NMDS) ordination via packages vegan and ggplot2 (Wickham 2016).

We identified significant indicator OTUs for each comparison group via the indicspecies package (De Caceres and Legendre 2009) using the ‘multipatt’ function with ‘IndVal’ setting, considering top indicator OTUs with a p value < 0.05. Using the package phyloseq (McMurdie and Holmes 2013), we compared across analysis groups and graphed for both digestive gland and hemolymph samples the percent relative abundance of all OUT’s that were deemed an indicator species for any of the three analysis groups.

### Bacterial isolation and *Yokenella* quantitative PCR (qPCR)

As a post-hoc procedure, we collected representative samples of hemolymph and digestive gland from four individuals for aerobic culture. These samples were processed by the UTCVM Bacteriology Lab. As a post-hoc analysis based on results from bacterial isolation and 16S sequencing, we further surveyed hemolymph and digestive gland DNA samples for *Yokenella* using the quantitative PCR (qPCR) to assess the association of disease with *Yokenella*. We followed the protocol described by Leis at el. (2023). Briefly, we used RxnReady™ Primer Pools for forward and reverse primers, PrimeTime™ qPCR probe with FAM - ZEN / Iowa Black™ FQ reporter dye - quencher combination and the PrimeTime™ Gene Expression Master Mix (Integrative DNA Technologies Inc., Coralville, IA, USA) with final concentrations matching the published protocol in a final reaction volume of 15 µL. We also used 217 bp synthetic gBlocks™ gene fragments (Integrative DNA Technologies Inc., Coralville, IA, USA) as listed in Leis et al. (2023) which included the 100bp target region of the *Yokenella pheS* gene as a positive control and for assay validation via standard curve. Standard curves were created from gBlocks™ reconstituted to 1 × 10^10^ copies/ µL with 10-fold serial dilutions down to 1 × 10^0^ copies/µL. We performed all assays in triplicate using the QuantStudio 6 (Applied Biosystems) at the University of Tennessee Institute for Agriculture (UTIA) Genomics Hub. Finally, we used Fisher exact test to evaluate the association between positive qPCR results for *Yokenella regensburgei* and lesions associated with gram-negative bacilli observed during histological assessment.

## Results

### Field sampling

We sampled control mussels at Kyles Ford (n =28) and Sycamore Island (n = 40) eight times in this study (data is S1 Table). During the summer of 2021, 32 control mussels died due to sedimentation induced smothering, caused by unusual high-water levels which made checking and cleaning silos impossible during that period. The remaining 68 control mussels did not present any signs of morbidity during the study period. Although most controls were sexually immature (n = 35), we identified 13 males and one female and were unable to identify the sex of 19 individuals due to their small size and absence of gonad in histological sections (data in S2 Table). At the time of silo deployment (May 2021) the median length of control mussels was 13.1 (ranging from 7.8 – 22.3mm, SD = 2). The median length of control mussels sampled from silos was 24mm (with a range of 16 – 34mm, SD = 4 mm) at Sycamore Island, and 24 mm (ranging from 16 – 26 mm, SD = 5 mm) at Kyles Ford during the duration of the experiment (May 2021 – November 2022).

We collected a total of 21 case mussels at Sycamore Island (n = 18), and Speers Ferry (n = 3; data in S1) over 24 surveys during October and November 2021 (n = 3), September – December 2022 (n = 9), and April and May 2023 (n = 9; data in S1 Table). We did not find case mussels at Kyles Ford. The sex ratio of cases was nearly equal, with 10 males, 9 females, and 2 hermaphrodites (both were male-dominant hermaphrodites with both ovarian and testicular gametogenic acini). The median size of case mussels was 81 mm (ranging from 60 – 109 mm, SD = 13mm) at Sycamore Island and 102 mm (ranging from 84 – 104 mm, SD = 11 mm) at Speers Ferry.

### Pathology

#### Controls

We examined 68 control mussels for pathology and did not observe any abnormalities grossly. Histologically, we frequently observed diffuse infiltrative hemocytosis (55/68, 81%; Fig 2B), hemocytic nodulation without histologically-evident infectious organisms (12/68, 18%), intracytoplasmic rickettsia like organisms within epithelium of digestive gland (29/67, 43%; Fig 2A), digenean trematode metacercaria (23/68, 34%), and nematodes (3/68, 4%; Table 1). Diffuse infiltrative hemocytosis affected the digestive gland (12/55, 18%), gonad (11/55, 20%), stomach (28/55, 52%), intestine (31/55, 46%), mantle (8/55, 12%), and gill (16/55, 23%). We observed hemocyte nodulation (Fig 2F), without associated necrosis or identified microorganisms (6/68), in the kidney (1/2, 50%) and intestinal lumen (1/2, 50%), and within the hemolymphatic spaces of the foot (1/4, 25%), gill (1/4, 25%), mantle (1/4, 25%), heart (1/4, 25%), and kidney (1/4, 25%; Table 1). We observed digenean trematode larvae, including encysted metacercaria in 22 control mussels and one that was too degenerated to determine larval stage in another mussel. In all instances metacercariae were in low number and hemocytic response was focal and mild. Hemocyte encapsulation (2/23, 9%), nodulation (10/23, 43%), or focal hemocytic infiltration (1/23, 5%) were associated with trematode larvae (Fig 2C), while 8/23 (24%) individuals showed no response and 2/23 (9%) showed multiple congruent responses (data is S3 Table). Nematodes were 10-20 µm diameter vermiform metazoa; one was within the digestive gland, and one was in the connective tissue at the base of the gill. Both had associated mild focal hemocytic nodulation with granulocytes predominating.

**Fig 2.**
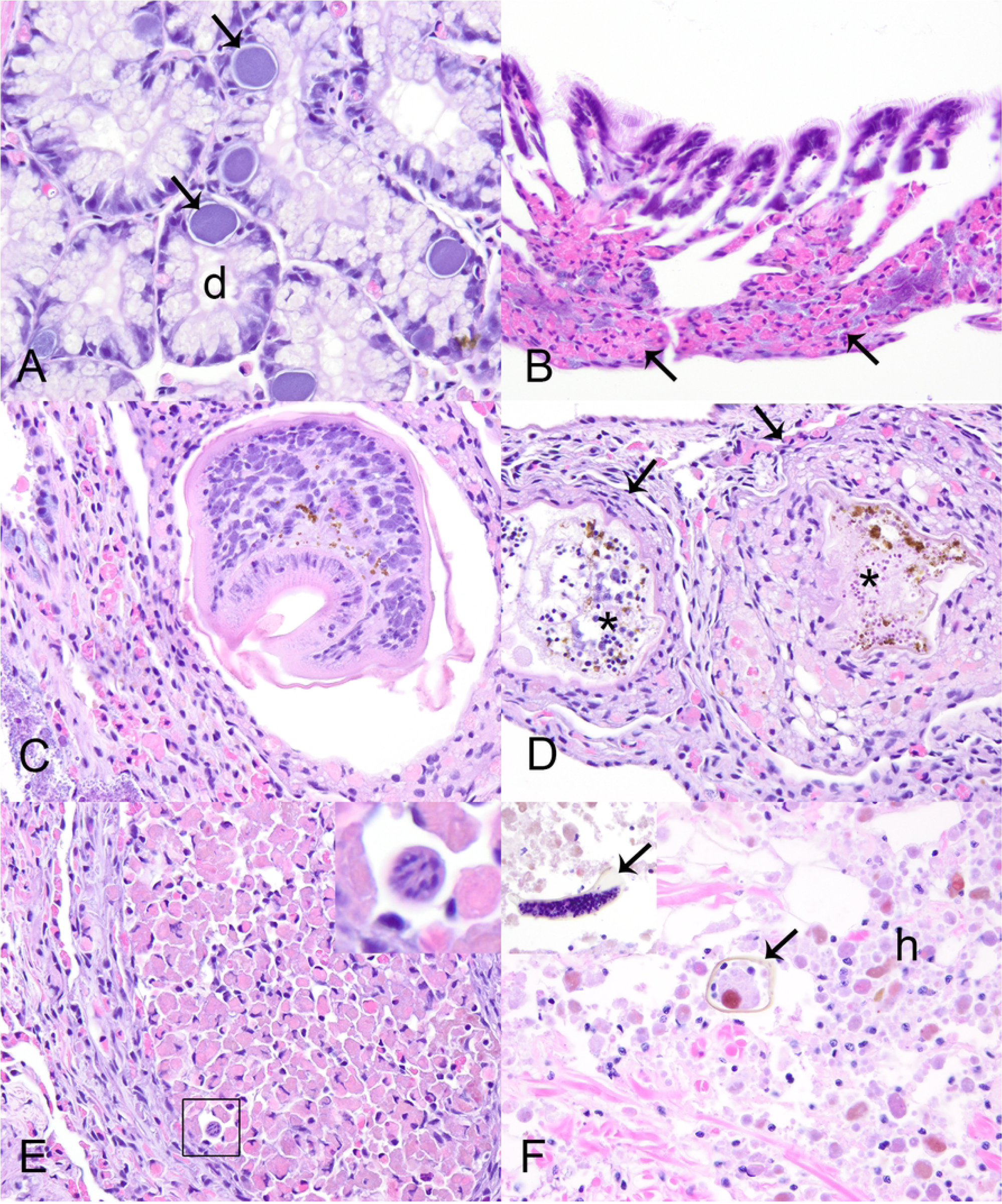
Microscopic lesions in control Pheasantshell (*Actinonaias pectorosa*). Microscopic lesions in control Pheasantshell (*Actinonaias pectorosa*) maintained in silos at sites impacted by seasonal mortality events in the Clinch River. A. Rickettsia-like organisms (arrows) within the epithelium of digestive gland tubules (d), H&E stain. B. Diffuse infiltrative hemocytosis within the gill connective tissue. Note the expansion of connective tissue at the base of lamellae by granular hemocytes (arrows), H&E stain. C. Encysted digenean trematode metacercaria within mixed hemocytic encapsulation, H&E stain. D. Encysted degenerate digenean trematode larvae (asterisks) bordered by mixed hemocytic encapsulation (arrows), H&E stain. E. Hemocytic encapsulation with primarily granulocytic hemocytes and containing a profile of a vermiform metazoan consistent with nematode (square, inset), H&E stain. F. Hemocytic nodulation centered on mite fragments (arrows, inset) as indicated by the refractile brown cuticle. Note the abundance of hemocytes with brown cytoplasmic pigment, H&E stain.

**Table 1.**
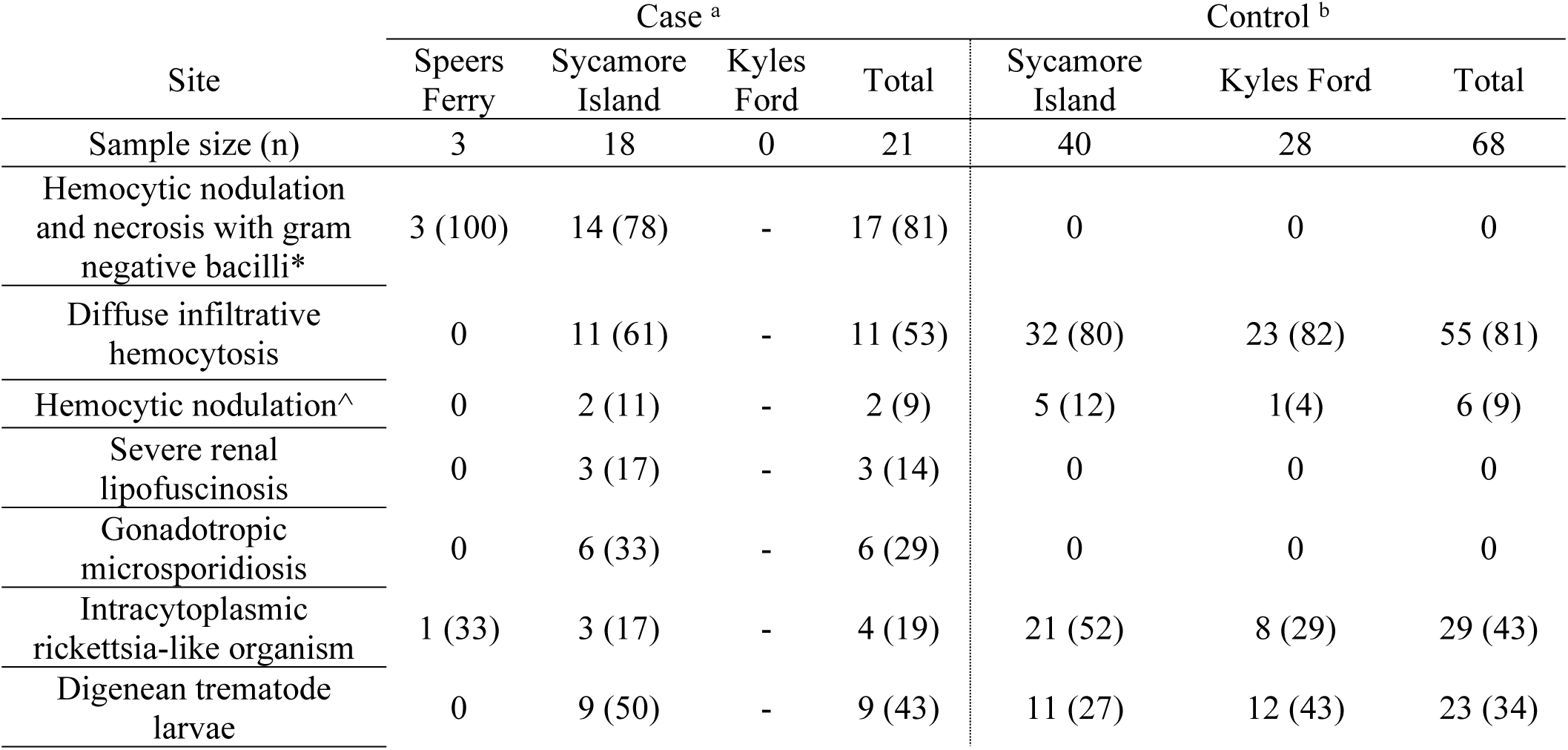
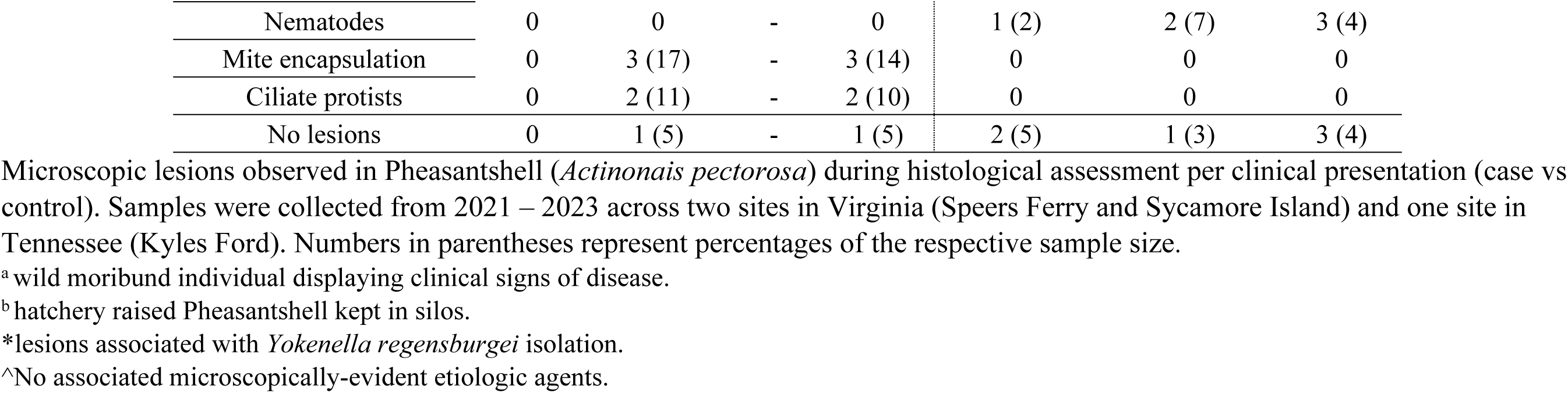
Microscopic lesions in Pheasantshell (*Actinonais pectorosa*) by clinical presentation and location (Virginia and Tennessee, USA)

#### Cases

We examined 21 case mussels for pathology. Grossly, we observed discrete multifocal dark brown to black streaks or patches in the marsupium of 3/7 (43%) gravid mussels (Fig 3A). One mussel had retraction of mantle at the posterior shell margin which covered anomalous shell, characterized by a fluid-filled, nacre-covered pocket (Figs. 3B-D). Microscopic examination of wet-mounted impressions of discolored marsupium revealed nonmoving glochidia with variably distinct internal tissue structure within brown-tinged mucus (S1 Figure, frame A). When similar preparations stained with modified Wright-Giemsa (Diff-Quik®) were examined microscopically, discolored areas contained brown pigment, agranular hemocytes, and glochidia lacking internal cellular detail, within a background of mucus, cellular debris, and bacilli (S1 Figure, frame B). In contrast, non-discolored areas had glochidia with preserved cellular detail, more mucus and ciliated cells, and fewer brown granules and bacteria (S1 Figure, frame C). Microscopic examination of the fluid within the anomalous shell failed to reveal parasitic agents. Cytological examination of modified Wright-Giemsa preparations of the shell fluid revealed the presence of crystalline and acellular amorphous debris, containing golden-brown granules, a variety of diatoms, bacilli, and cyanobacteria (Fig 3D).

**Fig 3.**
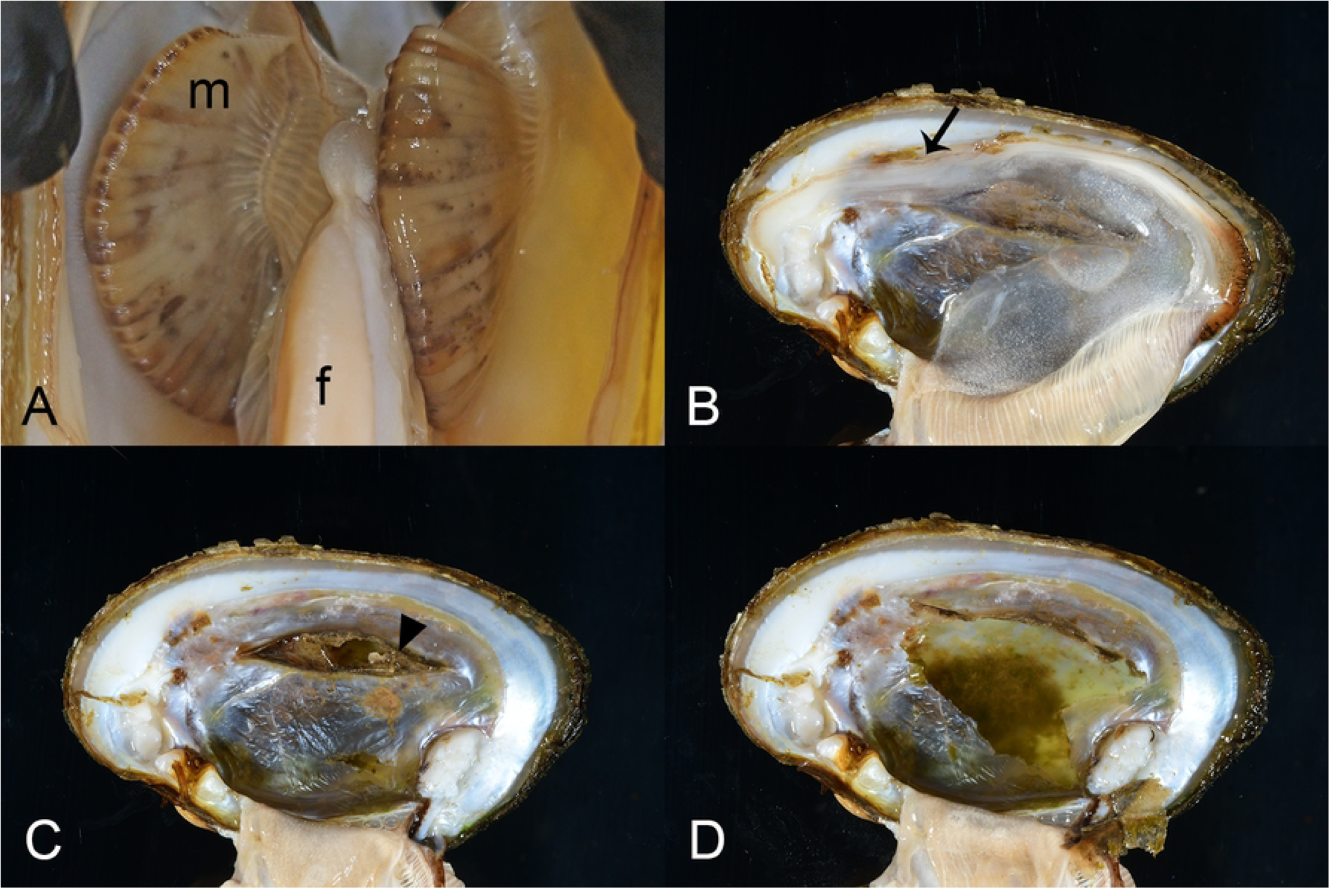
Gross lesions in case mussels. Gross lesions in case mussels (i.e., wild moribund Pheasantshell (*Actinonaias pectorosa*)) sampled at sites impacted by seasonal mortality events in the Clinch River. A. Gravid marsupium (m) discolored by brown foci; f – foot. B-D. Anomalous right shell valve characterized by recession of mantle at the posterior shell margin (B, arrow), which once removed, reveals an elevated thin layer of nacre with a hole (C, arrowhead) which covers a chamber filled with thin flocculent fluid containing green debris (D).

Histologically, 81% (17/21) of case mussels were affected by severe multi-organ hemocytic nodulation and necrosis associated with gram-negative bacilli (later identified as *Yokenella*), including all those with brown-black marsupium discoloration (Table 1). Affected mussels had multifocal aggregates of hemocytes within hemolymphatic spaces of many organs, most predominant in the connective tissue just beneath gastrointestinal tract (15/17, 88%; Figs. 4A-B) and integument surface of the visceral mass (15/17, 88%; Figs. 4C-D), as well as in digestive gland (9/13, 69%; Fig 4E), gonad (9/17, 53%), gill (15/16, 94%; Fig 4F), mantle (12/17, 71%), foot (12/17, 71%), kidney (11/16, 69%), heart (7/12, 58%; Fig 5A), and marsupium (4/6, 67%; Figs. 5B-C). Hemocytes in these aggregates were a mixture of predominantly agranular hemocyte (hyalinocytes) and fewer granulocytes. Centrally, they were necrotic as indicated by cytoplasmic hypereosinophilia, loss of cellular detail, and nuclear pyknosis or karyolysis. In all instances necrotic hemocytes were centered on intra- and extra-cellular slender long gram-negative bacilli in chains (Figs. 4B and 4D). Most mussels with multi-organ hemocytic nodulation and necrosis had some degree of epithelial cytoplasmic vacuolation associated with cell sloughing or erosion in the gastrointestinal tract or integument of the visceral mass. This change was often close to foci of hemocytic nodulation and sometimes contained similar bacteria (Figs. 4C and 5D). Hemocytic infiltration and necrosis occasionally involved the integument epithelium (3/17, 18%) or gastrointestinal epithelium or lumen (5/17, 29%; Fig 4A). In two mussels, hemocytic infiltrates also had features of encapsulation associated with the bacteria (Fig 5E). Additionally, some mussels with yokenellosis also had digestive gland epithelial necrosis (5/13, 38%; Fig 5F), as indicated by cellular dissociation, cytoplasmic hypereosinophilia, and loss of cell detail. Few mussels also had short gram-negative bacilli (n = 4), or large gram-positive bacilli (n = 4) within their tissues, but not localized to hemocytic infiltrate, and these were interpreted as perimortem opportunists.

**Fig 4.**
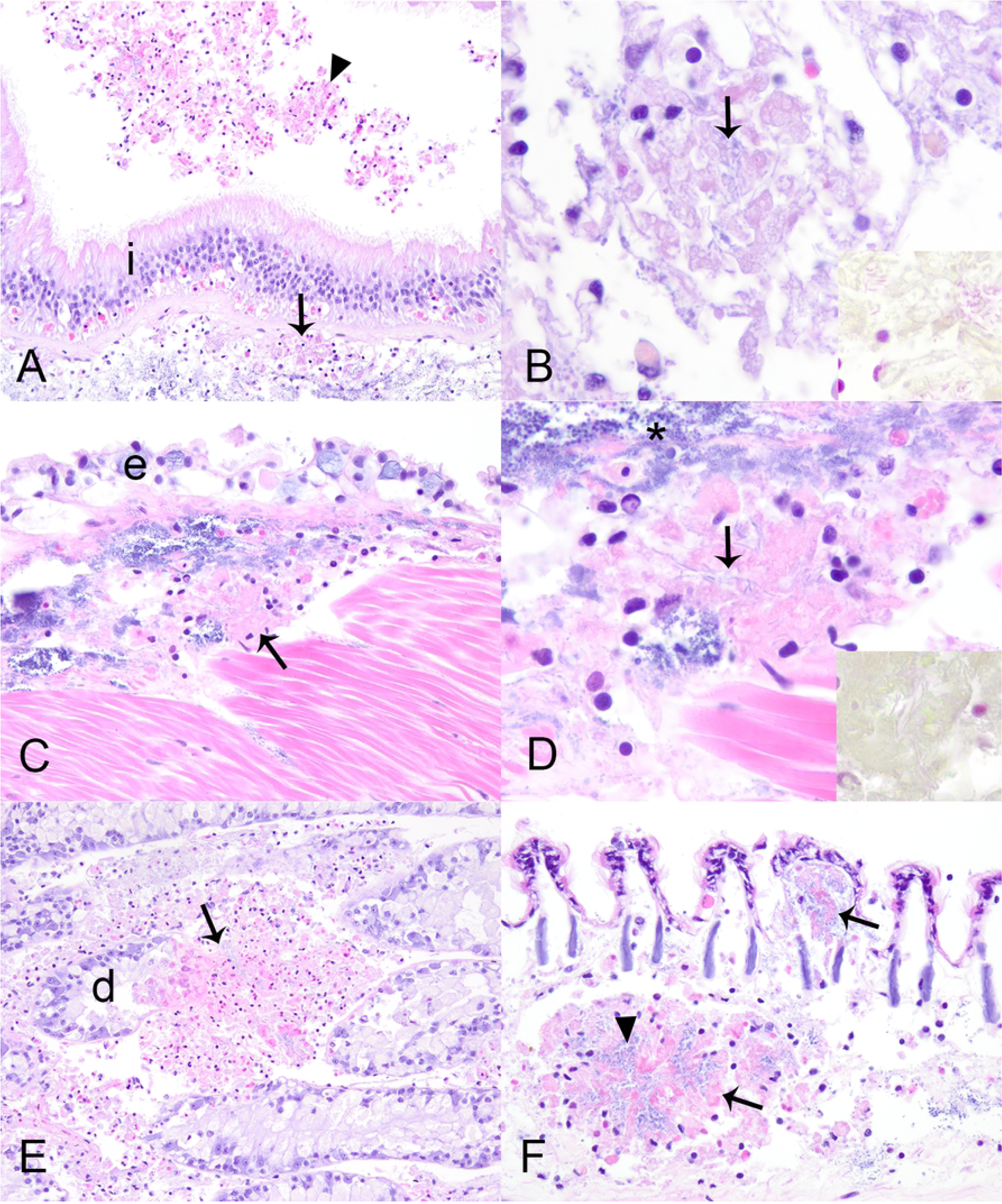
Microscopic lesions in case mussels. Microscopic lesions in case mussels (i.e., wild moribund Pheasantshell (*Actinonaias pectorosa*)) sampled at sites impacted by seasonal mortality events in the Clinch River. A. Hemocyte nodulation and necrosis within the submucosa (arrow) and lumen (arrowhead) of the intestine (i), H&E stain. B. Necrotic hemocytes center on slender bacilli (arrow) that are gram negative (inset, Hucker-Conn Gram stain), H&E stain. C. Hemocyte nodulation and necrosis beneath the epithelium (e) covering the visceral mass. Note vacuolation of the epithelium, H&E stain. D. Higher magnification of C, with necrotic hemocytes centered on slender bacilli (arrow) that are gram negative (inset, Hucker-Conn Gram stain), H&E stain. Basophilic granules (asterisk) are mineral concretions normally observed in connective tissues of freshwater mussels, H&E stain. E. Foci of hemocyte nodulation and necrosis (arrow) efface digestive gland acini (d), H&E stain. F. Foci of hemocyte nodulation and necrosis within the hemolymphatic spaces of the gill (arrows), similarly centered on bacilli (arrowhead), H&E stain.

**Fig 5.**
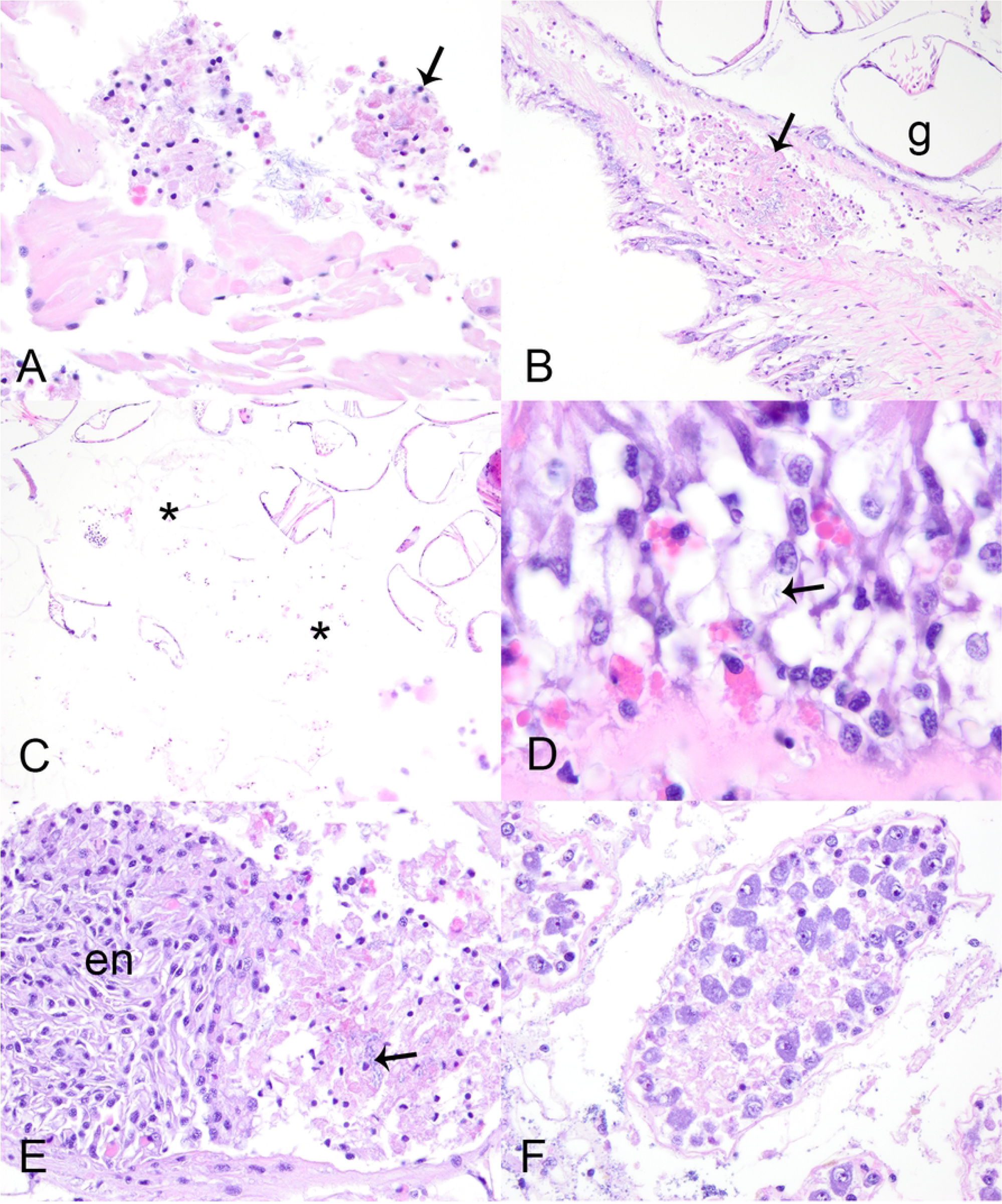
Microscopic lesions in case mussels. Microscopic lesions in case mussels (i.e., wild moribund Pheasantshell (*Actinonaias pectorosa*)) sampled at sites impacted by seasonal mortality events in the Clinch River. A. Luminal hemocytic aggregates (arrow) in the heart, H&E stain. B. Hemocytic nodulation and necrosis in the hemolymphatic space of the marsupium (arrow); g – glochidia, H&E stain. C. Glochidial necrosis (asterisks) within the marsupium, H&E stain. D. vacuolation of intestinal epithelium with intracytoplasmic bacilli (arrow), H&E stain. E. Early hemocytic encapsulation (en) adjacent to hemocytic nodulation and necrosis centered on bacilli (arrow), H&E stain. F. Necrosis of digestive gland acini, H&E stain.

Apart from the lesions attributed to yokenellosis, case mussels also had diffuse infiltrative hemocytosis (11/21, 53%), severe renal lipofuscinosis (3/21, 14%), gonadotropic microsporidiosis (6/21, 29%), intracytoplasmic rickettsia like organisms within epithelium of digestive gland (4/21, 19%; Fig 2A), digenean trematode metacercaria (9/21, 43%), mite encapsulation (3/21, 14%), and ciliates (2/21). Diffuse infiltrative hemocytosis affected the connective tissue around the stomach (11/20, 55%), intestine (7/21, 33%), mantle (1/21, 5%), ganglia (1/11, 9%), or gonad (1/21, 5%). We observed hemocyte nodulation (Fig 2F), without associated necrosis or identified microorganisms (2/21, 9%) in the gill (1/2) and adductor muscle (1/2). Trematode metacercariae had either hemocyte encapsulation (1/9, 11%), nodulation (3/9, 33%), or no cellular response to infection (5/9, 55%; data in S3 Table). Our chi-square test revealed an association between clinical presentation and diffuse infiltrative hemocytosis (p = 0.02), and no significant association with hemocytic nodulation, infection with digenean trematode, and intracytoplasmic rickettsia-like organism. Ciliated protists were located in the gill (1/2) and mantle (1/2). These were of one morphological type, consistent with mobile peritrchs: around 30µm diameter, dome to saucer shaped, with a dense lobed to horseshoe-shaped macronucleus, sparse somatic ciliature, and a refractile curvilinear structure (presumptive denticulate ring) bearing ciliary wreaths. In each case, they were in low numbers (<5) and not associated with pathology in the affected tissue.

Only four case mussels did not have lesions of yokenellosis. These mussels had severe gonadotropic microsporidiosis (n = 2, including the mussel with the anomalous shell), hemocytic encapsulation of degenerate mite fragments (n = 3; Fig 6B), and one had no lesions to explain morbidity (n = 1) except for hemocytic nodulation in the gills. One of the mussels with ovarian microsporidiosis had diffuse marked infiltrative hemocytosis of the visceral mass with intra-hemocytic microsporidia.

**Fig 6.**
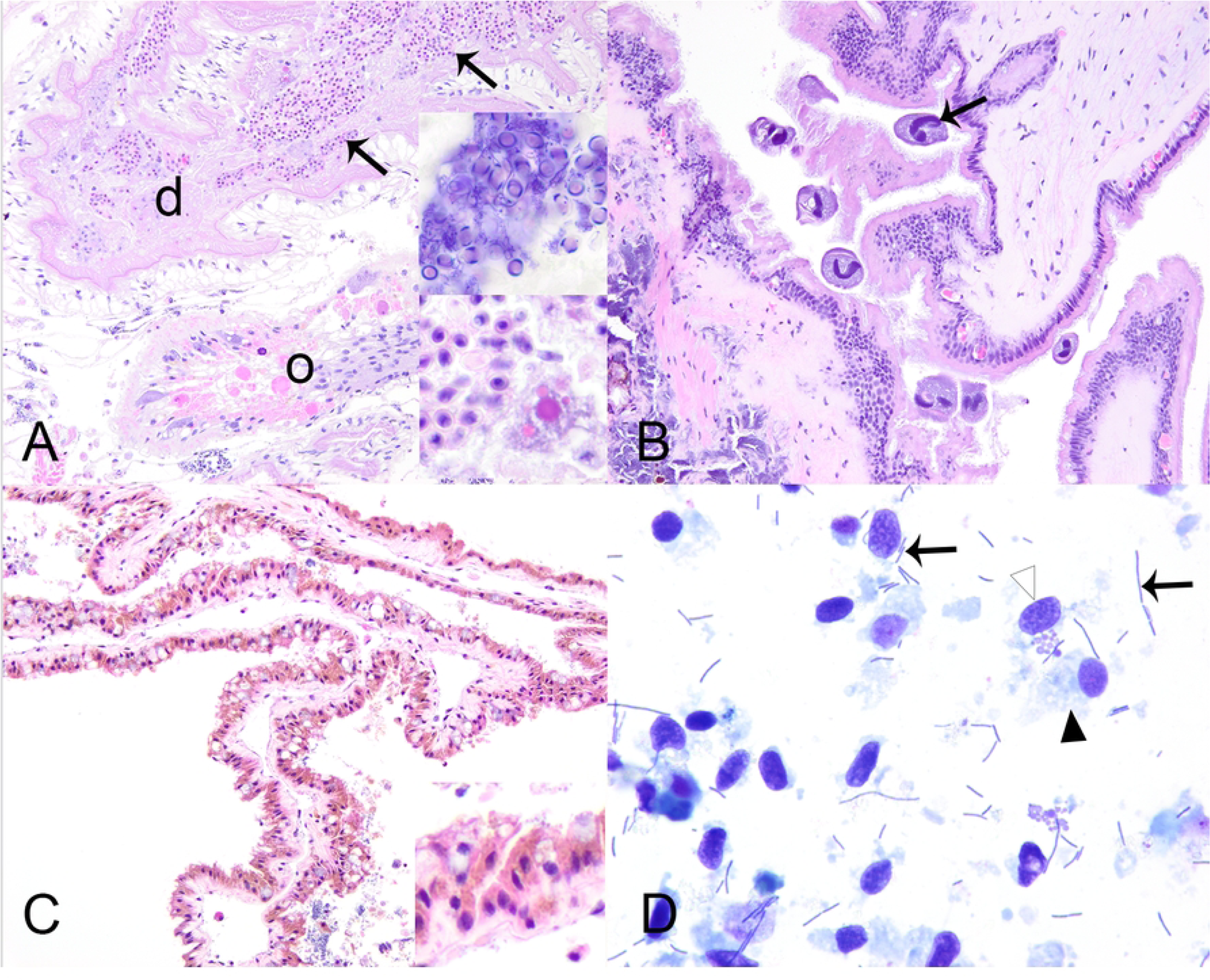
Microscopic lesions in case mussels. Microscopic lesions in case mussels (i.e., wild moribund Pheasantshell (*Actinonaias pectorosa*)) sampled at sites impacted by seasonal mortality events in the Clinch River. A. Gonadal duct (d) filled with mature oocytes containing numerous intracytoplasmic microsporidia (insets), H&E stain. The capsule of mature spores stain blue (gram positive; upper inset; Hucker-Conn Gram stain). Note depletion of ovarian acinus (o), H&E stain. B. Ciliate protist (arrow) infestation of mantle, H&E stain. C. Severe renal lipofuscinosis as indicated by the cytoplasm of nearly all nephron epithelium being distended with coarse brown granules (inset), H&E stain. D. Hemolymph smear containing numerous elongate bacilli in pairs and long chains (arrows) with predominance of agranular hemocytes (black arrowhead) over granular hemocytes (white arrowhead). Glochidial necrosis (asterisks) within the marsupium, Wright-Giemsa stain.

### Hemolymph cytology

Sufficient hemolymph was collected to allow cytological examination of 32 mussels, including 14 cases and 18 controls. One case mussel with yokenellosis produced very cloudy, white tinged, hemolymph. Hemocyte preservation was poor to fair in 26/32 samples and presented low cellularity in 7/32. We observed hemocyte clumping on 8/32 samples and diatoms in 14/32 samples. Lastly, we identified long unevenly stained bacilli, present as individuals, pairs, or filaments, in the hemolymph smears of 9/9 cases with yokenellosis (Fig 6D), in high (n = 7) or low (n = 2) numbers. Similar bacteria were not observed in the hemolymph smears of any cases without hemocytic nodulation and necrosis associated with gram-negative bacilli (n = 4), or control mussels (n = 18).

### Bacterial 16S amplicon high-throughput sequencing

We recovered 1,825,867 sequences representing 6,368 OTUs from 144 samples comprising 87 digestive glands (n = 68 control, 19 cases; 1,320,764 /5684), 46 hemolymph samples(n = 19 control, 13 cases; 412,106 / 1502). 4 community standards (92,754/ 187), 4 extraction controls (116 sequences / 48 OTUs), and 3 PCR blanks (127 / 16). After quality control removal of low abundance OTUs (<500 seqs), removal of low sequence yield samples (<200 seqs), and removal of control samples, there remained for analysis: 1,734,906 sequences representing 85 OTUs from 77 digestive glands (1251872/85), 13 hemolymph samples (39,1220/51), and 4 community standards (91814/16). Community standards were removed for all subsequent analyses. Hemolymph samples were included in general descriptive results and graphs for visual representation but were not used in statistical analysis.

Three analysis groups: case group with multi-organ hemocytic nodulation and necrosis associated with gram-negative bacilli (Case-Present, n = 11 digestive glands and 6 hemolymph samples), cases without hemocytic nodulation and necrosis associated with gram-negative bacilli (Case-Absent, n = 4 digestive glands and 1 hemolymph samples), and controls (Control-Absent, n = 62 digestive glands and 6 hemolymph samples). PERMANOVA and NMDS plots (Figs 7 and 8) revealed a significant effect of analysis group on digestive gland bacterial assemblage with only the pairwise comparison of Control-Absent vs Case-Present being significant for both Sorensen and Morisita-Horn indices (p = 0.001 and 0.019, respectively). Indicator species analysis revealed three OTUs indicative for Case-Present, seven OTUs for Case-Absent, and only one for the Control-Absent group (Table 2). The top two indicator OTUs for Case-Present mussels were OTU02 (a *Yokenella*/*Salmonella* sp.; p = 0.003) and OTU03 (*Aeromonas* sp.; p = 0.008). Relative abundance for these two OTUs was very high across both digestive gland and hemolymph samples from Case-Present and comparatively relatively low for both sample types of the other two groups. See data in S4 Table for entire list of indicator OTUs with average relative abundance listed by analysis group and sample type (hemolymph and digestive gland). Note amplicon sequences were only 250 bp which was not a large enough segment within the v4 region to distinguish *Yokenella* from *Salmonella* species, but we recorded OTU02 as *Yokenella* given supporting matched culture results.

**Fig 7.**
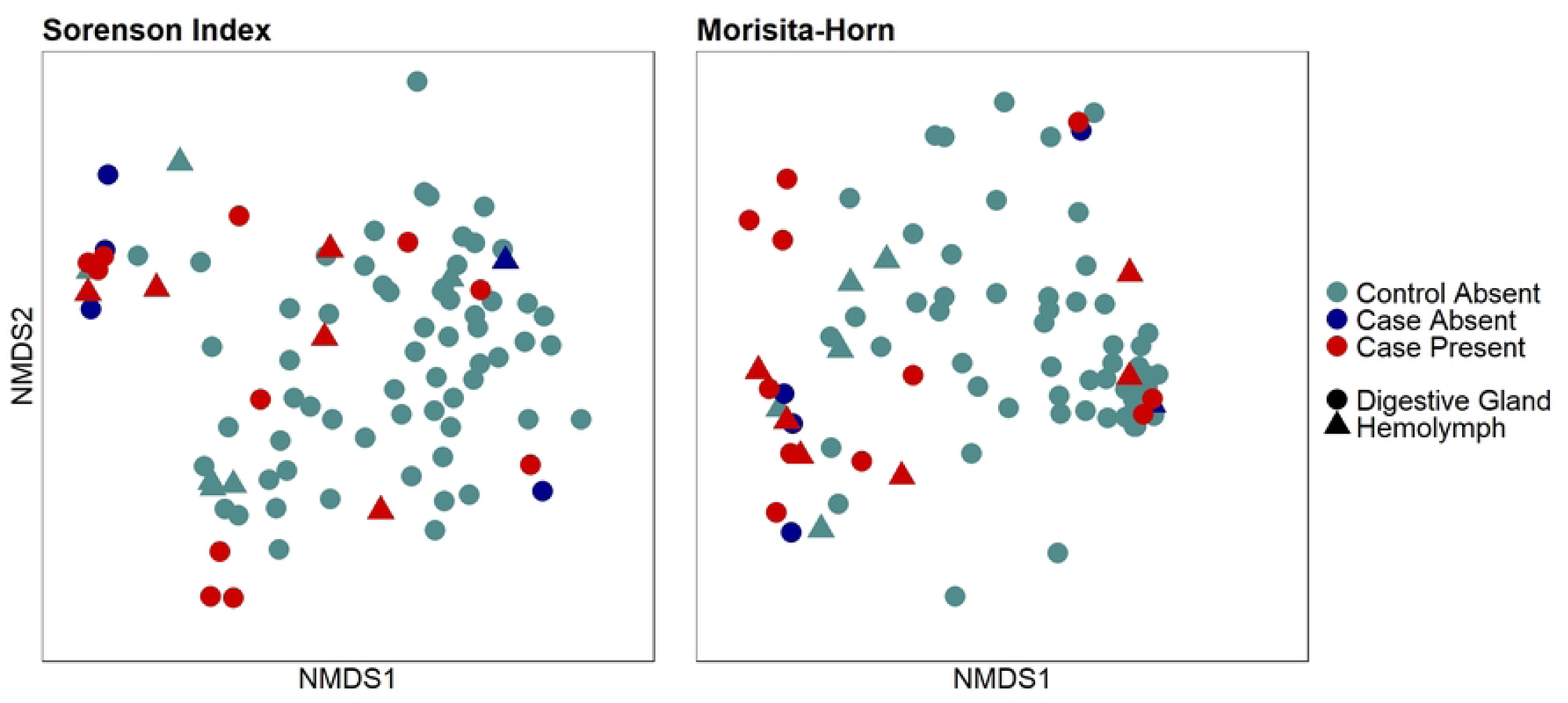
Non-Metric multidimensional scaling (NMDS) plot. A Non-Metric Multidimensional Scaling (NMDS) plot representing similarities and dissimilarities between analysis groups (Control-Absent; Case-Absent; Case-Present) and sample type (hemolymph and digestive gland). Each point represents an individual sample. Samples consisted of hemolymph and digestive gland tissue collected from Pheasantshell (*Actinonaias pectorosa*) sampled at sites impacted by seasonal mortality events in the Clinch River. Analysis groups include Control-Absent (individuals presenting no pathology; n = 62 DG and 6 HL), Case-Absent (individuals not presenting hemocytic nodulation and necrosis associated with gram-negative bacilli; n = 4 DG and 1 HL), and Case-Present (individuals presenting hemocytic nodulation and necrosis associated with gram-negative bacilli; n = 11 DG and 6 HL).

**Fig 8.**
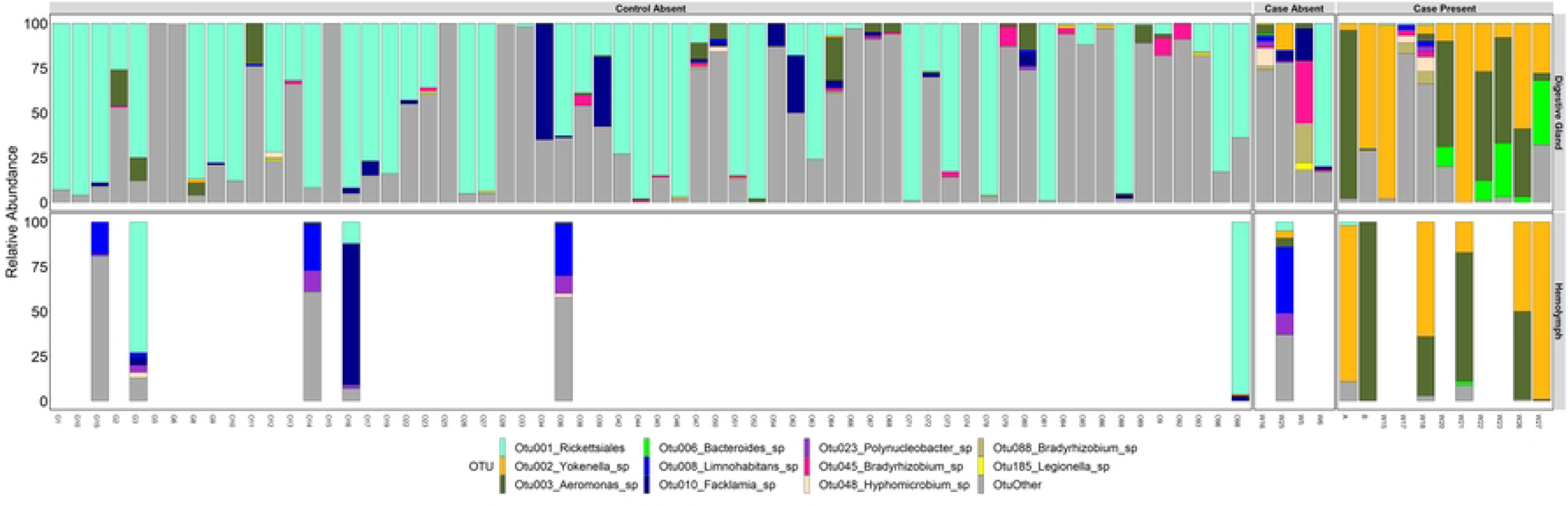
Relative abundance of operational taxonomic units (OTUs) per sample. Relative abundance of Operational Taxonomic Units (OTUs) per sample type and analysis group. Samples consisted of hemolymph (n=13) and digestive gland tissue (n=77) collected from Pheasantshell (*Actinonaias pectorosa*) sampled at sites impacted by seasonal mortality events in the Clinch River. Analysis groups include Control-Absent (individuals presenting no pathology; n = 62 DG and 6 HL), Case-Absent (individuals not presenting hemocytic nodulation and necrosis associated with gram-negative bacilli; n = 4 DG and 1 HL), and Case-Present (individuals presenting hemocytic nodulation and necrosis associated with gram-negative bacilli; n = 11 DG and 6 HL). Animal IDs are shown across the X axis and associated hemolymph and digestive gland samples linked.

**Table 2.**
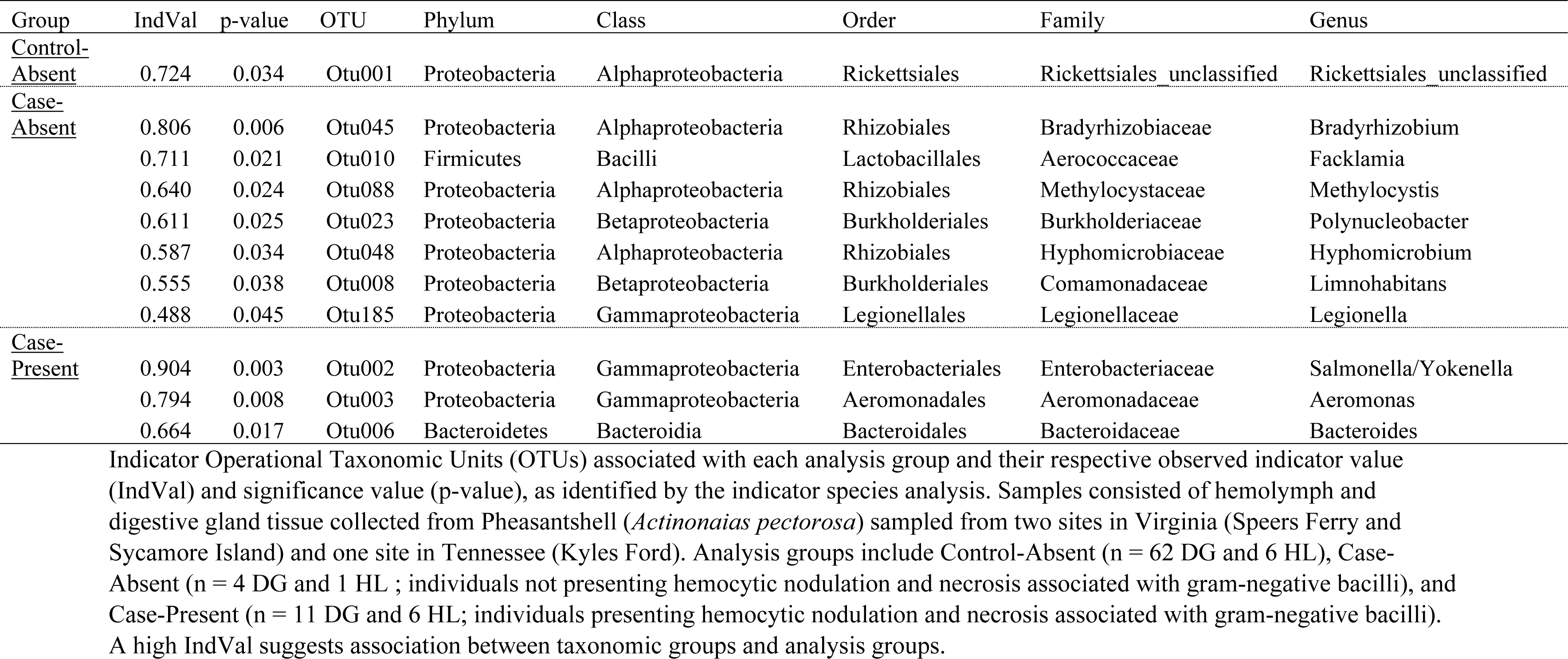
Indicator operational taxonomic units (OTUs) per analysis group.

### Bacterial isolation and *Yokenella* qPCR

Multiple bacteria were isolated from the digestive gland of four (IDs: W24, W25, W26, W27) and hemolymph of three (IDs: W24, W25, W26; Table 3) case mussels. Three out of four (W24, W26, W27) individuals had multi-organ hemocytic nodulation and necrosis associated with bacilli on histological examination and one (W25) had other lesions. *Yokenella regensburgei* was isolated from the hemolymph (1/3) and digestive gland (3/3) of all three individuals (W24, W26, W27) with multi-organ hemocytic nodulation and necrosis but was not isolated from the one individual (W25) without pathology associated with bacilli.

**Table 3.**
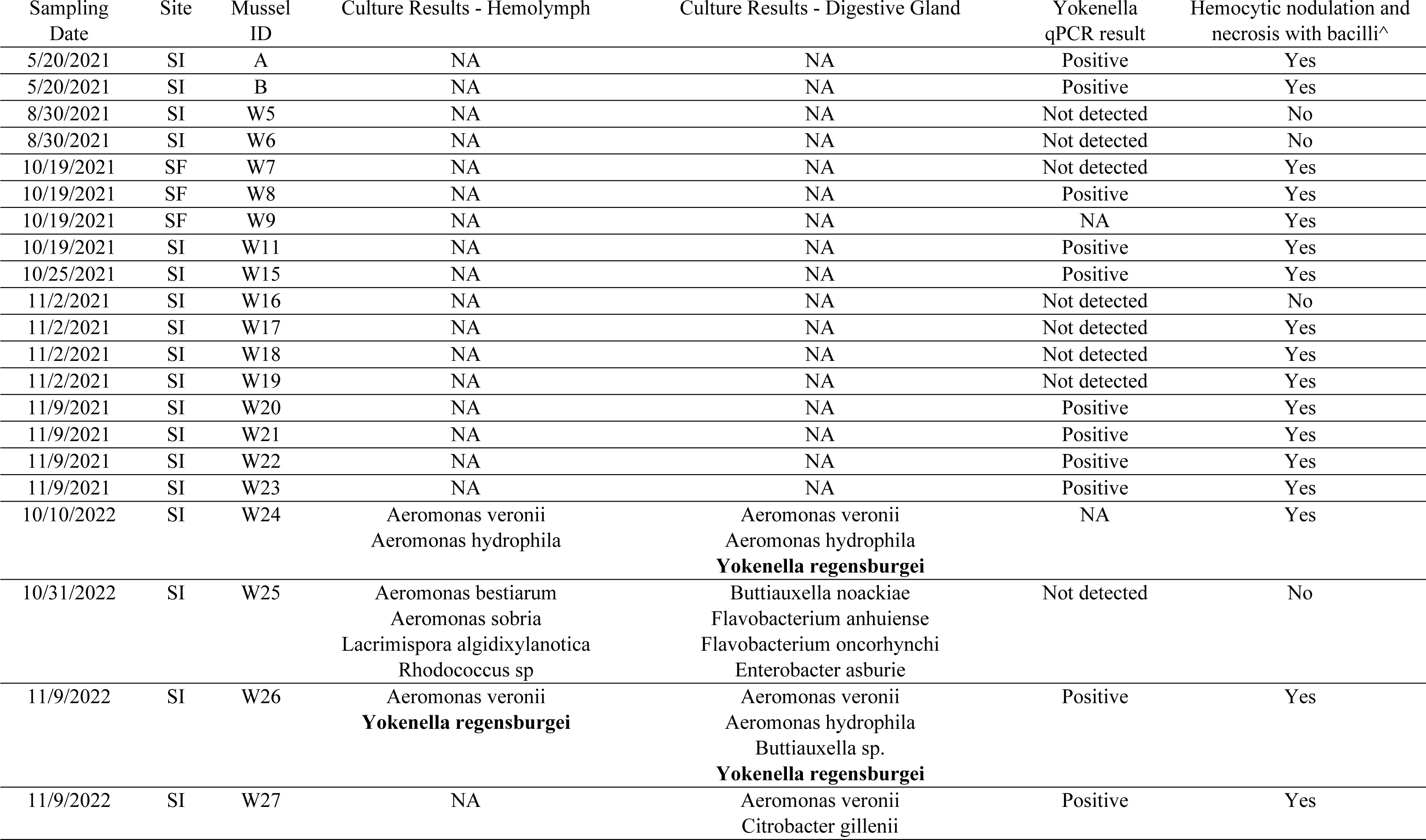
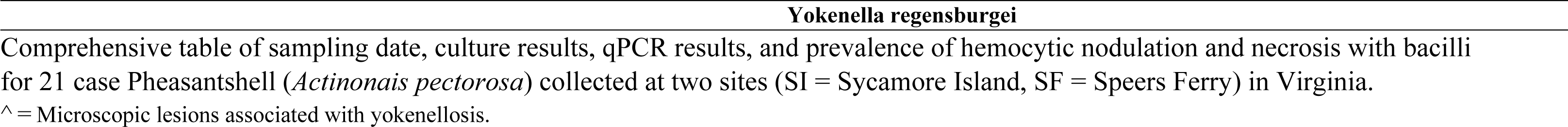
Summary of culture and qPCR results, along with hemocytic nodulation and necrosis prevalence in case pheasantshell (*Actinonais pectorosa*) from two study sites in Virginia.

We had 13 digestive gland DNA extraction samples from case mussels available for qPCR. We detected *Yokenella regensburgei* in 11/13 (85%) samples from cases and did not detect it in 17 randomly selected controls (Table 3). Although we were able to consistently amplify gBlock standards, our 10^0^-standard failed to amplify completely on all of our plates. Therefore, we considered our limit of quantification (LOQ) to be 10 copies/ µL. Severe multi-organ hemocytic nodulation and necrosis with gram-negative bacilli was associated with positive qPCR results for *Yokenella regensburgei* (Fisher exact test; p = 0.02).

## Discussion

In this study we investigated and described differences in pathology and microbiology between cases (i.e., wild moribund Pheasantshell) and controls (i.e., apparently-healthy hatchery-sourced Pheasantshell reared in silos) from a multi-year case-control study in the Clinch River. Most importantly, we use several methods to evaluate the role of bacteria in morbidity. Although our study cannot establish causation of mortality events, it provides the first robust evidence of a bacterium associated with pathology in wild freshwater mussels. Specifically, our results indicate that *Yokenella regensburgei* can be a pathogenic bacterium in free-living Pheasantshell at our study sites, confirming that at least part of the mortality observed in the Clinch River can be attributed to infectious disease.

In our study, we found that 81% of cases were affected by a characteristic lesion (multi-organ hemocytic nodulation and necrosis), that consistently had intra- and extra-cellular slender long gram-negative bacilli in chains typical of Enterobacteriaceae. *Yokenella regensburgei* was consistently isolated from mussels with this lesion when bacterial culture was undertaken, but not from those lacking this lesion, indicating the identity of intralesional bacteria. Additionally, relative abundance of a *Yokenella/Salmonella* sp. OTU was very high across both digestive gland and hemolymph samples from cases with hemocytic nodulation and necrosis compared to controls or cases without this lesion (<5% for both across both sample types). We also successfully detected *Y. regensburgei* in 69% of individuals with these lesions via qPCR compared to 0 detections in individuals without. *Yokenella regensburgei*, identified through both culture and genetic sequencing, has previously been detected in low prevalence in tissue homogenate of moribund Ebonyshell (*Fusconaia ebena*) and in high prevalence in the hemolymph of moribund Pheasantshell from mass mortality events in the Tennessee River, Alabama, and in the Clinch River, Tennessee, respectively (Starliper et al. 2011; Leis 2019; Richard et al. 2021). Despite previous studies reporting the presence of *Y. regensburgei* in freshwater mussels, the present study is the first to link the pathogen to pathology in the infected host. The lesion descriptions provided by this study can serve as a resource for those investigating freshwater mussel mortality. Specifically, yokenellosis is a primary differential for foci of hemocytic nodulation and necrosis containing bacilli. Moreover, the associated lesions are of sufficient severity and physiological significance to explain mortality in infected hosts. Taken together, these findings indicate *Yokenella regensburgei* can be a significant pathogen in Pheasantshell. While its role in causing mortality would ultimately require confirmation via experimental infection trials, meeting Koch’s postulates will present a variety of challenges given the difficulties in recreating a conducive aquatic environment and maintaining healthy but susceptible hosts in a laboratory setting. Multifactorial causal models that account for enabling and predisposing factors, such as environmental interactions and microbial dysbiosis, are likely more appropriate for establishing etiology in infectious invertebrate diseases (Rothman 1976; Vonaesch et al. 2018).

To the authors’ knowledge, bacterial diseases have not yet been described in wild freshwater mussels (Knowles et al., 2023). This finding is remarkable because bacterial diseases in bivalves more broadly are largely limited to aquaculture, particularly hatchery-reared juvenile stages, facilitated by various artificial conditions (Paillard and Borrego 2004; Travers et al. 2015). Multi-organ hemocytic nodulation and necrosis, the primary lesion of moribund mussels, is consistent with lesions observed in septic diseases of other mollusks (Terio et al. 2018; De Vico and Carella 2012). Findings indicating chronic infection or debilitation such as emaciation or mantle retraction were absent in cases. The presence of a developing encapsulation-type hemocytic response in two mussels suggests infection may not necessarily be acute and rapidly-progressing, but this should be confirmed by clinical observations. If bacterial infection was an opportunistic terminal event, pathological conditions predisposing to infection were not evident in this study.

Curiously, previous histological evaluations of Pheasantshell from the Clinch River did not reveal similar lesions. Henley et al. (2019) examined 10 visibly unhealthy mussels (gaping shell, poor ability to retract foot) from Kyles Ford in October 2016, and described putative cocci, but not associated lesions, and a variety of hemocytic infiltrates (termed aggregations, vesicular occlusions, or granulocytomas) not clearly associated with necrosis or gram-negative bacilli. Knowles et al. (2022) examined 8 moribund mussels from Kyle’s Ford and Speer’s Ferry in October 2018, 4 of which were infected with a gonadotropic microsporidian, *Knowlespora clinchi* (Bojko 2022). In that study, the observed site-specific mortality patterns did not correlate with the prevalence of *K. clinchi*, suggesting that mortality did not appear to be the result of *K. clinchi*. Moreover, lesions similar to those attributed to yokenellosis in our study were not identified (personal communication, S. Knowles). The failure of these studies to document hemocytic nodulation and necrosis with bacilli may reflect differences in timing of sampling relative to disease or mortality occurrence or may indicate that yokenellosis is one of multiple problems leading to Pheasantshell mortality, supported by a minority of our case mussels having pathological conditions other than yokenellosis, such as microsporidiosis, or shell disease. Consistent with this view, multi-pathogen mortality events are increasingly recognized in aquatic invertebrates, particularly where driven by a complex interplay of environmental stressors (Bower et al. 1994; Austin and Austin 2012; Carella et.al 2023). Our surveys in Kyle’s Ford did not yield moribund mussels, limiting comparisons to these studies. However, Leis et al. (2019) isolated *Y. regensburgei* from mussels during a mortality event in Kyle’s Ford, November 2018, but histology was not undertaken by that study and the pathological significance of *Y. regensburgei* was undetermined at that time. This scenario emphasizes the value of a paired sampling approach, where multiple tests can be completed on a single mussel in parallel to facilitate interpretation of test results.

Hemolymph assessment has been used to evaluate the health of freshwater mussels and shows value as a non-lethal method for disease surveillance and diagnostics in freshwater mussels (Gustafson et al. 2005a,b; Burkhard et al. 2009). We identified bacilli in all hemolymph smears of case mussels with yokenellosis (Fig 6D). Bacterial 16S sequencing and culture results from hemolymph were also informative, highlighting differences in abundance of *Yokenella* and other bacteria taxa between cases and controls. Changes in hemolymph color from transparent to a cloudy, white tinged color may be a gross indicator of bacterial infection. In this case, microscopic assessment should be conducted to confirm the diagnosis. Overall, hemolymph assessment can be a non-lethal tool to study and diagnose yokenellosis in endangered wild mussels.

*Yokenella regensburgei* is a gram-negative bacillus and the only species in its genus. In humans, it is an opportunistic pathogen that rarely causes disease, infecting a wide variety of tissues in almost exclusively immunosuppressed individuals (Lo et al. 2011). Apart from freshwater mussels, its only documentation in animal disease involves a recent report of fatal pneumonia and septicemia in a population of farmed American alligators (*Alligator Mississippians*) in Louisiana (Balamayooran et al. 2022). The source of infection or route of transmission were not identified in our study or by Balamayooran et. al (2002). Some infected mussels had lesions involving gill, integument, alimentary tract, or kidney, providing a route for bacterial entry, or shedding in the water column. Conditions which would predispose mussels to infection were similarly unclear. There were a variety of other lesions (not associated with bacteria) observed at similar prevalences in case and control mussels, contributing to a broader understanding of “background” (non-mortality-associated) lesions in mussels (Burcham et al 2023; Knowles et al. 2023). Gonadotropic microsporidia were unique to case mussels, potentially reflecting immunosuppression, or simply predilection for larger (sexually mature) mussels.

Our control group never developed disease while wild Pheasantshell did. We hypothesize that differential disease occurrence may be explained by superior health status and resiliency of controls, smaller mussels being less vulnerable to infection, silos protecting from exposure to etiologic agents in the substrate, or low prevalence of disease more broadly. Although our control group was genetically related to wild mussels from the Clinch River, they were raised in a hatchery, in a stable environment with controlled climate, flow and abundant food, and consequential robust health status could explain resistance to infection once translocated. This possibility is supported by our data where control mussels did not show signs of disease despite some with low abundance of the *Yokenella* sp. OTU. It is also possible to this cohort did not have long enough exposure to unknown environmental stressors required to facilitate disease. In addition, keeping mussels in silos with limited contact with the substrate may have reduced exposure to etiologic agents in the benthos through burying or pedal feeding. Finally, there is still little known regarding the epidemiology of the mortality events observed in the Clinch River; the failure to observe disease in control mussels may simply indicate disease prevalence is <1% (and therefore may not be detected when examining only 68 individuals), or that disease preferentially affects mussels of a larger size. We were unable to harvest apparently healthy size matched wild Pheasantshell for our control group due to regulatory constraints. Given the growing uncertainty around the representativeness of hatchery-derived or silo-maintained controls, it is important that access to apparently healthy wild mussels is expediated for those undertaking diagnostic investigations of unusual mortality events in mussels.

Microbial assemblage showed an overall significant difference between analysis groups. This was driven by the overwhelming abundance of *Yokenella* sp. and *Aeromonas* sp. OTUs that represented the majority of sequences in both hemolymph and digestive gland yokenellosis samples. *Aeromonas* spp. are commonly found in aquatic environments and are generally accepted as opportunistic invaders (Wiklund et al. 1998; Harikrishnan et al. 2005; Austin and Austin 2012). Like with *Yokenella*, *Aeromonas* was in very low relative abundance from cases without yokenellosis (Fig 8; Case-Absent) and controls (Fig 8; Control-Absent). Our results are supported by other studies that have observed co-occurrence of *Aeromonas* and *Yokenella* in moribund mussels (Leis et al. 2019, Richard et al. 2021, Leis et al. 2023). Species of this genus may be facultative pathogens to freshwater mussels that proliferate in the right conditions, such as in the presence of already damaged tissue or weak/stressed host, commonly co-infecting mussels with yokenellosis. Cooperative polymicrobial infections are a common feature among *Aeromonas* sp. infection in other species and have similarly been documented in vibriosis of marine bivalves (Lemire et al. 2015; Fernández-Bravo and Figueras 2020; Cantillo et al. 2023). However, a third bacterial OTU, a *Bacteroides* sp, was also preferentially abundant in these animals and significantly associated with this group (Case-Present) via indicator species analysis. The abundance of this OTU within the yokenellosis group was still highly variable and likely the reason behind why NDMS plots do not show a complete separate grouping of yokenellosis cases from others. It is unclear if this *Bacteroides* sp. plays a role in pathology. This genus is commonly associated with gut flora, is an obligatory anaerobe, and has been proposed to be used as an indicator of water quality and mammalian fecal contamination (Teixeira et al. 2020), raising questions around wastewater exposure as a potential source of *Y. regensburgei* in this population. Alternatively, one *Bacteroides* species has been cultured from river sediment in Japan independent from known animal host contamination (Ismaelia et al. 2018), supporting that this particular OTU may be normal river flora. The source of this OTU and its relative abundance in the environment should therefore be further explored alongside *Yokenella*.

Interestingly, in contrast to cases with yokenellosis, controls and cases without yokenellosis had a dominant digestive gland OTU belonging to the order Rickettsiales. This OTU was especially abundant in controls representing nearly half (45%) of all recovered sequences in those digestive glands, and was the only OTU significantly associated with this group via indicator species analysis. Rickettsia-like organisms have been sporadically reported via histological examination of the digestive gland of mussels previously (Mioduchowska et al. 2020; Burcham et al. 2023) and could represent an important commensal in healthy tissue. This rickettsial OTU was also found with high relative abundance in the hemolymph of controls while lower in hemolymph of both case groups (Case-Present, Case-Absent) and warrants further investigation for use as a potential indicator of mussel health. Finally, indicator species analysis also revealed presence of a *Limnohabitans* sp. in the digestive gland of cases without yokenellosis (Case-Absent). This genus is well known to naturally inhabit freshwater systems (Kasalický et al., 2013) and has been identified as abundant on lesioned vs healthy skin of aquatic salamanders (hellbenders, *Cryptobranchus alleganiensis*) which have a range that overlaps with several species of freshwater mussels in the eastern and midwestern United States (Hernández-Gómez et al. 2017). Like *Bacteroides*, the role of *Limnohabitans* in the health of freshwater mussels should be investigated further.

Overall, our study provides baseline information and description of normal and abnormal cell and tissue anatomy of freshwater mussels, which is generally lacking (Knowles et al. 2023; McElwain and Bullard 2014). Most importantly, we provide a comprehensive pathological description of lesions associated with *Y. regensburgei* and address its pathogenicity (Leis et al. 2023). Although our study does not explain the cause of these infections, it confirms that mussels at two of our study sites are ultimately dying from an infectious disease and that *Y. regensburgei* can be pathogenic in free-living mussels. Our study reveals the importance of pairing pathology with other diagnostic tests when investigating invertebrate disease and mortality. It also highlights the need for epidemiological investigations that identify environmental and host factors conducive to bacterial disease epidemics and for experimental trials with *Y. regensburgei* to further investigate its pathogenicity.

## Acknowledgments

We thank the One Health Initiative at The University of Tennessee, Knoxville, the University of Tennessee College of Veterinary Medicine Comparative and Experimental Medicine Program, the McClung Museum of Natural History and Culture, and Morris Animal Foundation (D21ZO-802) for funding this project. We thank Jordan Richard, Rose Agbalog, and Andrew Henderson for conceptual guidance and Justin Wolbert, Devin Henever, Cameron Mitchell, and Merrie Urban for field assistance. We thank Sujata Argawal for technical assistance with qPCR and Veronica Brown for technical assistance with 16S library preparation and sequencing. We that the staff of the Aquatic Wildlife Conservation Center of Virginia Department of Wildlife Resources for rearing and providing control mussels deployed to silos. We also thank the Clinical Pathology, Histology, and Bacteriology laboratory staff for technical assistance. We also thank Emily Ford for editing and formatting support and Dr. Rebecca Bergee for statistical analysis support.

## Supporting information

**S1 Figure. Cytological examination of gill tissue.** Cytological examination of tissues of case mussels (i.e., wild moribund Pheasantshell (*Actinonaias pectorosa*)) sampled at sites impacted by seasonal mortality events in the Clinch River. A. Wet-mounted impression of discolored marsupium with brown-tinged mucus and glochidia having poorly defined tissues (inset). B. Impression smear of discolored marsupium with abundant brown-granules (arrows) and agranular hemocytes (arrowhead) within a background of mucus containing abundant bacilli (inset right) and glochidia with poorly discerned tissues (inset left), modified Wright-Giemsa stain. C. Impression smear of normal-colored area of the same marsupium as B, comparatively more mucus (m), intact ciliated epithelium (arrow) and hemocytes (arrowhead), and glochidia with discernable inner tissues (inset), modified Wright-Giemsa stain. D. Smear of fluid within the chamber of an anomalous shell consisting of amorphous acellular and crystalline debris, with scattered brown granules (black arrowhead), diatoms (arrow, inset upper), and cyanobacteria (white arrowhead, inset lower), modified Wright-Giemsa stain.

**S1 Table. Pheasantshell (*Actinonaias pectorosa*) Sampled by Clinical Presentation between Virginia and Tennessee Sites**. Total number of Pheasantshell (*Actinonaias pectorosa*) sampled by clinical presentation (case vs control) from two sites in Virginia (Speers Ferry and Sycamore Island) and one site in Tennessee (Kyles Ford).

**S2 Table. Pheasantshell (*Actinonais pectorosa*) Sampled by Sex and Clinical Presentation across Virginia and Tennessee Sites.** Pheasantshell (*Actinonais pectorosa*) sampled by sex and clinical presentation from two sites in Virginia (Speers Ferry and Sycamore Island) and one site in Tennessee (Kyles Ford).

**S3 Table. Trematode Prevalence in Pheasantshell (*Actinonais pectorosa*) Sampled across Virginia and Tennessee Sites.** Total number of Pheasantshell with trematode larvae per clinical presentation (case and control) identified in Pheasantshell (*Actinonaias pectorosa*) sampled from 2021 – 2023 across from two sites in Virginia (Speers Ferry and Sycamore Island) and one site in Tennessee (Kyles Ford), and total trematode count per site. We also identified trematode life stage and hemocyte response per clinical presentation across all three sites.

**S4 Table. Indicator Operational Taxonomic Units (OTUs) Relative Abundance in Pheasantshell (*Actinonaias pectorosa*) Hemolymph and Digestive Gland Tissues Sampled across Virginia and Tennessee Sites.** Average relative abundance of indicator Operational Taxonomic Units (OTUs) per sample type and analysis-group. Samples consisted of hemolymph and digestive gland tissue collected from Pheasantshell (*Actinonaias pectorosa*) sampled from the Clinch River at two sites in Tennessee and Virginia. Analysis groups include Control-Absent (n = 62 DG and 6 HL), Case-Absent (n = 4 DG and 1 HL ; individuals not presenting hemocytic nodulation and necrosis associated with gram-negative bacilli), and Case-Present (n = 11 DG and 6 HL; individuals presenting hemocytic nodulation and necrosis associated with gram-negative bacilli). Values in bold highlight the highest relative abundance per group.

